# Telomere-to-telomere genome assembly of matsutake (*Tricholoma matsutake*)

**DOI:** 10.1101/2022.07.25.501483

**Authors:** Hiroyuki Kurokochi, Naoyuki Tajima, Mitsuhiko P. Sato, Kazutoshi Yoshitake, Shuichi Asakawa, Sachiko Isobe, Kenta Shirasawa

## Abstract

Here, we report the first telomere-to-telomere genome assembly of matsutake (*Tricholoma matsutake*), which consists of 13 chromosomes (spanning 160.7 Mb) and a 76 kb circular mitochondrial genome. The chromosome sequences were supported with telomeric repeats at the ends. GC-rich regions are located at the middle of the chromosomes and are enriched with long interspersed nuclear elements (LINEs). Repetitive sequences including long-terminal repeats (LTRs) and LINEs occupy 71.7% of the genome. A total of 28,322 potential protein-coding genes and 324 tRNA genes were predicted. Sequence and structure variant analysis revealed 2,322,349 single nucleotide polymorphisms and 102,831 insertions and deletions, 0.6% of which disrupted gene structure and function and were therefore classified as deleterious mutations. As many as 683 copies of the LTR retrotransposon *MarY1* were detected in the matsutake genome, 91 of which were inserted in gene sequences. In addition, 187 sequence variations were found in the mitochondrial genome. The genomic data reported in this study would serve as a great reference for exploring the genetics and genomics of matsutake in the future, and the information gained would ultimately facilitate the conservation of this vulnerable genetic resource.

## Introduction

Matsutake (*Tricholoma matsutake* [S. Ito et Imai] Singer), belonging to the phylum Basidiomycota, is an ectomycorrhizal fungus that coexists with Pinaceae and Fagaceae trees in a symbiotic association^1,2^. In the field, two spores of matsutake fuse together and grow to form a “shiro”, which is a symbiotic entity formed between matsutake and its host tree. One shiro produces a number of sporocarps during the growing season. The sporocarp of matsutake has been considered as one of the most valuable components of traditional Japanese cuisine since ancient times, as mentioned in Manyo-shu (a series of books for Japanese poetry compiled around 700 AD in Japan), owing to its pleasant aroma, which is largely attributed to 1-octen-3-ol (also known as matsutakeol)^3,4^; however, sporocarps are non-culturable. In 2019, the International Union for Conservation of Nature (IUCN) categorized matsutake as vulnerable. The production of sporocarps has drastically decreased in recent years^5^ because of the deterioration of its growing environment. To understand the life cycle and life history of matsutake, safeguarding its production and conservation is necessary, which requires genomic analysis.

Four assemblies of the matsutake genome are currently available in a public DNA database^6,7^. However, the sequences are highly fragmented because contigs are enormous in number (2,545 to 88,884) and short (N50 length = 2.9 to 320.9 kb), thus providing insufficient genome coverage. Moreover, because retrotransposons such as *MarY1* span ~6 kb in length and are dispersed throughout the matsutake genome^8^, a full-length genome assembly may not be achieved with short-read and error-prone long-read sequencing technologies, both of which were employed to construct the four genome assemblies. Another reason why achieving a full-length genome assembly might be difficult is the diploid nature of the matsutake genome; it is difficult for symbiotic fungi to produce mononuclear hyphae (monokaryon) with haploid genomes. Unlike symbiotic fungi, saprophytic fungi produce mononuclear hyphae, and therefore can be easily sequenced using short-read and/or error-prone long-read technologies to obtain long contiguous genome assemblies.

Recently, the development of high-fidelity long-read (HiFi) technology (PacBio, Menlo Park, CA, USA) enabled the establishment of complete gapless assemblies of the human genome at the telomere-to-telomere level^9^, in which a single contig corresponds to a single chromosome. In this study, we applied the HiFi technology to address the complexity of the matsutake genome. Using this technology, we determined the total chromosome number of matsutake, which is consistent with the results of the few cytogenetics studies conducted to date^10^. Overall, this study represents a milestone in the cytogenetics-, genetics-, and genomics-focused research on matsutake mushroom.

## Materials and methods

### Fungus material and DNA extraction

Two sporocarps, which were probably ramets derived from a single shiro (radius > 2 m) that has been generating sporocarps for more than 20 years^11^, were collected from Ina, Nagano, Japan. The sporocarps were flash-frozen in liquid nitrogen, dried under vacuum, and then stored at room temperature until needed for DNA extraction.

Genomic DNA was extracted from the dried stipes using the cetyltrimethylammonium bromide (CTAB) method^12^. The concentration of the extracted DNA was measured using the Qubit dsDNA BR assay kit (Thermo Fisher Scientific, Waltham, MA, USA), and DNA fragment length was evaluated by agarose gel electrophoresis with Pippin Pulse (Sage Science, Beverly, MA, USA).

### DNA sequencing

Genomic DNA was subjected to HiFi SMRTbell library construction using the SMRTbell Express Template Prep Kit 2.0 (PacBio), according to the manufacturer’s instructions, with a minor modification. Because the genomic DNA was degraded, the DNA shearing step recommended in the protocol was skipped. The resultant DNA was fractionated with BluePippin (Sage Science) to eliminate fragments less than 10 kb in size. The DNA libraries prepared from the two sporocarps were indexed with unique barcode adapters, and sequenced on a single SMRT cell 8M on the Sequel IIe system (PacBio).

### Genome assembly and gene annotation

Using the HiFi reads obtained from the Sequel IIe system (PacBio), the genome size of matsutake was estimated with GCE^13^, based on *k*-mer frequency (*k* = 21) calculated with Jellyfish^14^ (version 2.3.0). The reads were assembled using hifiasm^15^ (version 0.16.1), with default parameters. Assembly completeness was evaluated with Benchmarking Universal Single-Copy Orthologs (BUSCO)^16^ (version 5.2.2; default parameters) using lineage dataset agaricales_odb10 (eukaryota, 2020-08-05). Telomere sequences containing repeats of a 6 bp motif (5’-CCCTAA-3’) were searched by BLASTN^17^ (version 2.2.26), with an E-value cutoff of 1E-20. Nuclear genes were predicted with Funannotate (https://doi.org/10.5281/zenodo.2604804) (version 1.8.9) using RNA-Seq reads downloaded from the NCBI nucleotide database (accession number: SRR485866). Mitochondrial genes were predicted with Artemis^18^, in accordance with the gene sequences reported in previous mitochondrial genome assemblies (accession number: NC_028135). The predicted genes were functionally annotated with emapper^19^ (version 2.1.6; search option: mmseqs) implemented in EggNOG^20^, and with DIAMOND^21^ (version 2.0.13; more sensitive mode) search against the UniProtKB^22^ database. Simultaneously, gene sequences reported in the previous genome assembly, Trima3^6^, were mapped on to the current assembly with Liftoff^23^ (version 1.6.3; parameter: -polish). Repetitive sequences in the assembly were identified with RepeatMasker (https://www.repeatmasker.org) (version 4.1.2; parameters: -poly and -xsmall) using repeat sequences registered in Repbase^24^ and a *de novo* repeat library built with RepeatModeler (https://www.repeatmasker.org) (version 2.0.2a; default parameters). Sequences showing similarity to *MarY1* (accession number: AB028236; 6047 bp) and its long terminal repeats (LTRs; 426 bp) were searched by BLASTN^17^.

### Sequence variant analysis

Single nucleotide polymorphisms (SNPs) and insertions and deletions (indels) were detected with genome sequence reads obtained from NCBI database (accession number: PRJNA726361). Low-quality bases and adapter sequences were removed with PRINSEQ^25^ (version 0.20.4) and fastx_clipper (parameter, -a AGATCGGAAGAGC), respectively, in the FASTX-Toolkit (version 0.0.14; http://hannonlab.cshl.edu/fastx_toolkit). The remaining high-quality reads were mapped on to the current assembly with Bowtie2^26^ (version 2.3.5.1; parameters: --local -I 100 -X 1000), and sequence variants were detected using the mpileup and call commands of BCFtools^27^ (version 1.9). High-confidence variants were selected with VCFtools^28^ (version 0.1.12b) using the following parameters: minimum read depth ≥ 8 (--minDP 8); minimum variant quality = 999 (--minQ 999); maximum missing data < 0.5 (--max-missing 0.5); and minor allele frequency ≥ 0.05 (--maf 0.05). Insertions of *MarY1* were detected with PTEMD^29^ (version 1.03). Effects of nucleotide sequence variations on gene function were estimated with SNPeff^30^ (version 4.3t).

Four matsutake genome assemblies downloaded from the NCBI database (accession numbers: BDDP01, Tricma30605_assembly01; PKSN02, ASM293902v2; QMFF01, ASM331463v1; WIUY01, Trima3) were aligned against the genome assembly generated in this study using Minimap2^31^ (version 2.24; parameter: -cx asm20), with a mapping-quality cutoff of 60. The positions of genes, repeats, and genome alignments were compared using the intersection command in BEDtools^32^ (version 2.27.0; default parameters).

## Results

### DNA sequencing, data analysis, and genome assembly

Genomic DNA was extracted from two dried sporocarps (samples A and B) of matsutake. The amount of DNA extracted from each sample (9 µg) was sufficient for library construction; however, because of degradation (Supplementary Figure S1), the extracted DNA was used for library preparation without shearing. The resultant libraries were sequenced on a SMRT Cell 8M to obtained 9.5 Gb (sample A) and 7.8 Gb (sample B) data, with N50 lengths of 11 kb (sample A) and 10 kb (sample B). The *k*-mer analysis detected two peaks (Supplementary Figure S2), indicating that the haploid genome size of matsutake was 149 Mb and the level of heterozygosity was high. The sequence reads of each sample were assembled separately to obtain two sets of contigs: 182 contigs (165.5 Mb) for sample A, and 146 contigs (162.9 Mb) for sample B. Among these, 15 contigs (160.7 Mb) of sample A and 12 contigs (159.2 Mb) of sample B, all of which were >1 Mb in length, were selected for further analysis.

Next, we searched for the telomeric motif, (CCCTAA)n, in the contigs (Figure 1). In sample A, the telomeric motif was found at both ends of nine contigs (A1, A2, A6, A8, A11, A12, A13, A14, and A16) and at one end of five contigs (A3, A4, A7, A10, and A15). In sample B, the motif was found at both ends of nine contigs (B1, B2, B3, B4, B5, B7, B8, B9, and B11) and at one end of three contigs (B6, B12, and B13). The average length of the telomeric sequence was 129 bp (21 repeats), and ranged from 66 bp (11 repeats) to 168 bp (28 repeats).

**Figure 1.**
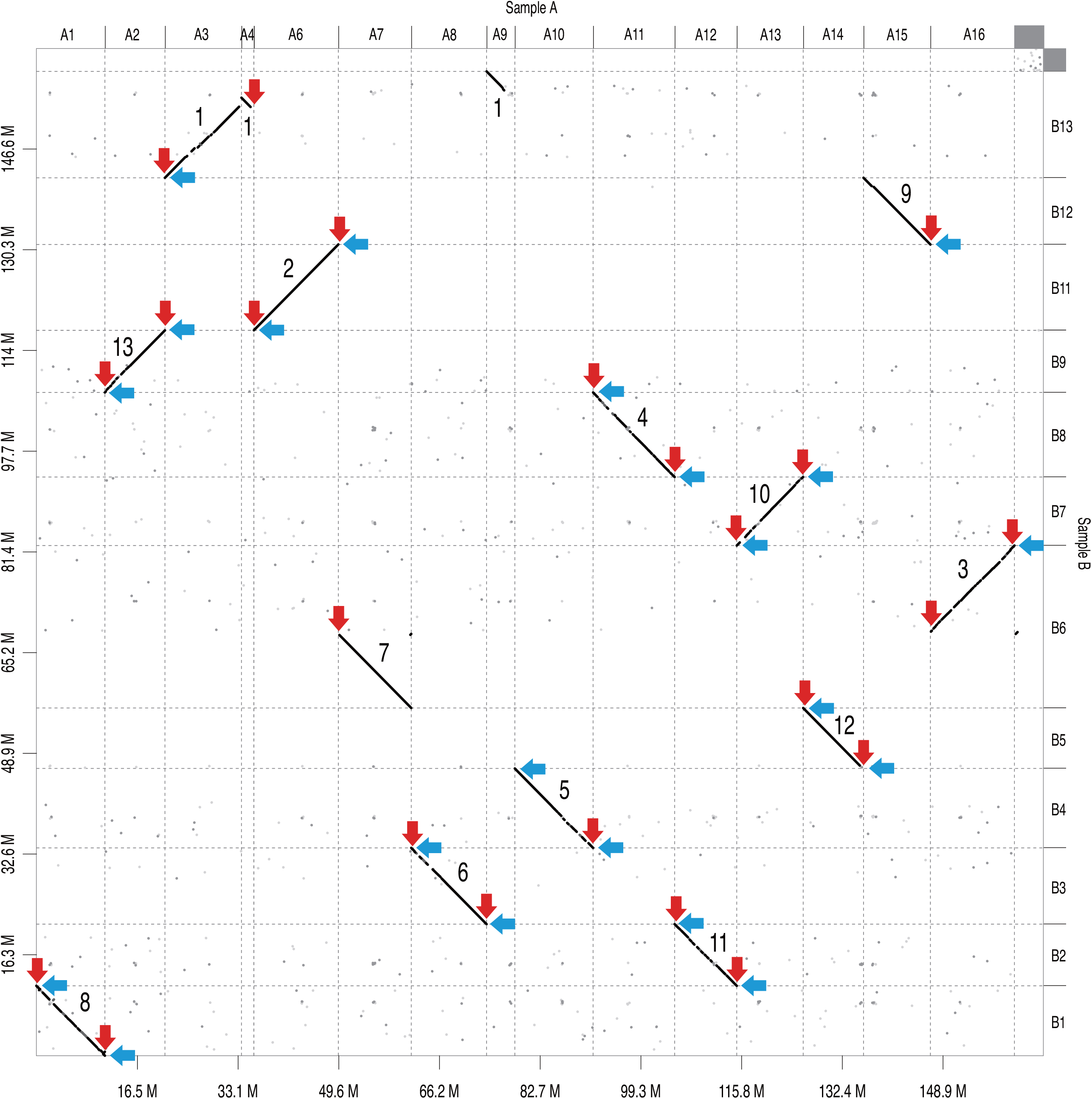
Comparative map of contigs of samples A and B. Dots indicate sequences similar between the two samples. Red and blue arrows indicate telomeric motifs detected at the ends of contigs of samples A and B, respectively. Numbers in the plot indicate chromosome numbers in the final assembly (TMA_r1.0). Contigs A5 and B10 are lacked because of the short sequence length (<1 Mb).

Comparison of the two sets of genome assemblies revealed 10 pairs of perfectly aligned contigs (A1-B1, A2-B9, A6-B11, A8-B3, A10-B4, A11-B8, A12-B2, A13-B7, A14-B5, and A15-B12) (Figure 1). Three contigs of sample A (A3, A4, and A9) covered the entire sequence of one contig of sample B (B13). Furthermore, two contigs of sample A (A7 and A16) corresponded to one contig of sample B (B6). Thus, we concluded that contigs A3, A4, and A9 were unassembled, and contig B6 was misassembled. Therefore, we joined contigs A3, A4, and A9 with 100 Ns to establish a single contig, and left contigs A7 and A16 as separate. Finally, 13 contigs spanning 160.7 Mb were obtained, of which 11 contigs possessed telomeric motifs at both ends, while two contigs were supported with the telomeric motif at either end. The 13 contigs represented 94.2% complete BUSCOs. The final assembly was designated as TMA_r1.0, and the contigs were named TMA_r1.0ch01 to TMA_r1.0ch13 in order of decreasing sequence length (Figure 1, Table 1). The GC content was ca. 45% over the entire genome, with one peak (~55%) in each chromosome, except chromosome 1, which showed two peaks (Figure 2). In addition, we identified a 76,067 bp circular contig, which represented the mitochondrial genome of matsutake (Tma1.0mito).

**Table 1.** Statistics of the matsutake genome assembly

| Chromosome | Sequence length (bp) | No. of genes | Contigs of sample A | Contigs of sample B |
| --- | --- | --- | --- | --- |
| Tma1.0ch01 | 19,249,005 | 3,124 | A3, A4, A9 | B13 |
| Tma1.0ch02 | 13,874,649 | 2,147 | A6 | B11 |
| Tma1.0ch03 | 13,809,479 | 2,888 | A16 | B6 (bottom) |
| Tma1.0ch04 | 13,409,878 | 2,584 | A11 | B8 |
| Tma1.0ch05 | 12,860,286 | 2,423 | A10 | B4 |
| Tma1.0ch06 | 12,376,747 | 2,372 | A8 | B3 |
| Tma1.0ch07 | 11,987,682 | 2,101 | A7 | B6 (top) |
| Tma1.0ch08 | 11,264,506 | 2,027 | A1 | B1 |
| Tma1.0ch09 | 10,996,241 | 1,862 | A15 | B12 |
| Tma1.0ch10 | 10,915,969 | 1,886 | A13 | B7 |
| Tma1.0ch11 | 10,203,996 | 1,724 | A12 | B2 |
| Tma1.0ch12 | 9,912,917 | 1,714 | A14 | B5 |
| Tma1.0ch13 | 9,869,253 | 1,794 | A2 | B9 |
| **Total** | **160,730,608** | **28,646** |  |  |

**Figure 2.**
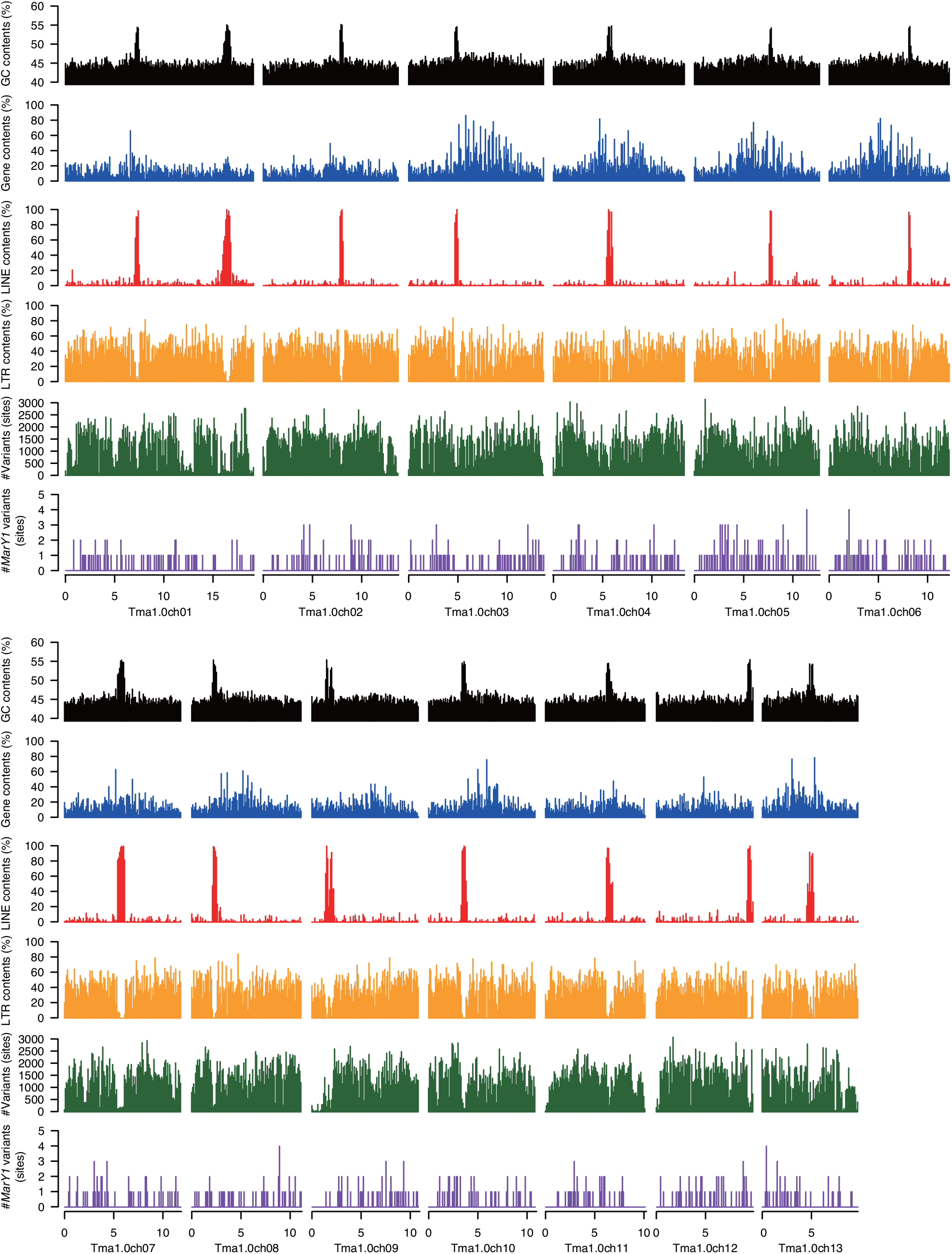
Structures and components of the matsutake genome. Bars indicates the GC content (black) and numbers of genes (blue), LINEs (red), and LTRs (orange) within a 100 kb window. Green and purple bars indicate the number of sequence variants (SNPs and indels) and number of *MarY1* insertions, respectively.

### Repetitive sequence analysis

Repetitive sequences occupied a total physical distance of 115.2 Mb (71.7%) in the genome assembly (TMA_r1.0; 160.7 Mb). Nine major types of repeats were identified in varying proportions (Table 2). The dominant repeat types in the chromosome sequences were LTRs (69.2 Mb) and long interspersed nuclear elements (LINEs; 5.9 Mb). LINEs were predominant in regions with high GC content in all chromosomes, whereas LTR retrotransposons were predominant in regions with low GC content (Figure 2). Repeat sequences unavailable in public databases totaled 16.2 Mb. The *MarY1* LTR, which has been extensively studied to date, and its terminal repeats were present as 683 and 3,240 copies, respectively, across all 13 chromosomes.

**Table 2.** Repetitive sequences in the matsutake genome

| Type of repetitive sequence | Copy number | Length (bp) | Proportion of genome (%) |
| --- | --- | --- | --- |
| SINEs | 304 | 51,368 | 0.0 |
| LINEs | 7,377 | 9,428,495 | 5.9 |
| LTR elements | 60,502 | 69,150,848 | 43.0 |
| DNA transposons | 10,805 | 7,760,391 | 4.8 |
| Small RNA | 362 | 70,121 | 0.0 |
| Satellites | 81 | 17,133 | 0.0 |
| Simple repeats | 9,262 | 422,530 | 0.3 |
| Low complexity | 970 | 56,370 | 0.0 |
| Unclassified | 68,440 | 26,094,240 | 16.2 |

### Gene prediction and annotation

RNA-Seq reads were mapped on to the genome assembly of matsutake (TMA_r1.0), with a mapping rate of 93.8%. Based on the sequence alignment, TMA_r1.0 was predicted to contain a total of 28,646 genes including 28,322 protein-coding genes and 324 tRNA genes (Table 1). These predicted genes possessed 90.2% complete BUSCOs. The mitochondrial genome was predicted to contain a total of 28 protein-coding genes, including 26 tRNA genes and 2 rRNA genes.

Additionally, sequence alignment revealed that of the 23,068 genes predicted in the previous assembly (Trima3), 22,326 were represented in the current assembly (TMA_r1.0). Comparison of the genome positions of the two gene sets indicated that 12,761 of the 28,646 genes predicted in TMA_r1.0 overlapped with 13,082 of the 22,326 genes in Trima3. The remaining 15,885 genes (= 28,646 – 12,761) were unique to TMA_r1.0.

### Comparative analysis of the current and previous genome assemblies of matsutake

The TMA_r1.0 genome assembly was compared with the four publicly available matsutake genome assemblies, Tricma30605_assembly01, ASM293902v2, ASM331463v1, and Trima3. Sequence coverage in GC-rich regions was mostly low in the four assemblies (Supplementary Figure S3). The Tricma30605_assembly01 covered the longest part of TMA_r1.0 (79.5%) among the four assemblies, followed by Trima3 (78.9%), ASM331463v1 (75.9%), and ASM293902v2 (67.9%). When genomic positions of the alignments with the four assemblies were merged, 94.4% of the TMA_r1.0 was covered by at least one of the four assemblies, while the remaining 5.6% was not covered by any assembly.

### Sequence variants in divergent matsutake lines

Whole-genome resequencing data of 14 matsutake lines were obtained from a public DNA database. High-quality reads (3.9 Gb per sample) were mapped on to TMA_r1.0, with an average mapping rate of 96.5%, except one sample (TM_NH), which showed a mapping rate of 73.3%. Totals of 2,322,349 SNPs and 102,831 indels were identified in the 13 chromosomes (Figure 2). The most prominent variant type was intergenic mutations (2,106,014, 86.8%) followed by missense mutations

(148,235, 6.1%) and synonymous mutations (99,674, 4.1%) (Supplementary Table S1). The number of deleterious variations, which could disrupt gene structure and function, was 13,479 (0.6%); these were categorized as high-impact variants.

*MarY1* insertions were detected in all 13 chromosomes at 747 positions across the 14 lines (Figure 2). The number of *MarY1* insertions per line ranged from 67 in EF to 135 in W2. Among the 747 positions, 91 were located within the gene coding sequence, and 556 were located in upstream and downstream regions of genes.

In the mitochondrial genome, a total of 90 SNPs and 97 indels were identified across all 14 lines, although no *MarY1* insertion was detected. Deleterious mutations were found in three mitochondrial genes, *orf123, orf290*, and *cox1*.

## Discussion

This study presents the telomere-to-telomere genome sequence of matsutake comprising 13 chromosomes (Figure 1, Table 1). To assemble the matsutake genome, we not only considered the telomeric repeat motif but also identified the centromeric regions and sequenced two independent samples. GC-rich regions were found at a single position in all chromosomes, except chromosome 1, which had two GC-rich regions (Figure 2). Interestingly, the GC-rich regions were enriched with LINEs but devoid of LTRs (Figure 2). Together, these observations suggest that GC-rich regions represent centromeres, and that chromosome 1 is a dicentric chromosome formed by the telomeric fusion of two chromosomes. We also compared the genome assemblies generated from two independent data sets (samples A and B) (Figure 1). Consequently, it was possible to identify a misassembled region and an unassembled region (Table 1), which led to the establishment of a telomere-to-telomere genome assembly. To the best of our knowledge, haploid chromosome number of matsutake (n = 7) has been reported in only one study to date^10^. Constructing a telomere-to-telomere assembly could serve as an alternative to karyotyping for determining the chromosome number of a species, for which no chromosome information is available.

The telomere-to-telomere genome assembly generated in this study spans a physical distance pf 160.7 Mb and covers the entire genome of matsutake. The genome size of matsutake is larger than that of other mushroom species^6,7^ because of the high proportion of repetitive sequences (Table 2)^33^. Owing to its high content of repetitive sequences (Table 2) and high heterozygosity (Supplementary Figure S2), the matsutake genome could not be fully sequenced with short-read and error-prone long-read sequencing technologies. The HiFi sequencing technology (~10 kb read length) employed in this study likely helped overcome the problem posed by repetitive sequences, such as *MarY1* (~6 kb), thus enabling the construction of the telomere-to-telomere genome assembly. Owing to the long contigs and high genome coverage, 28,646 genes were predicted in the matsutake genome. Of these genes, 15,885 had not been represented in the previous assembly (Trima3).

The genome sequences and predicted genes could help us understand the ecophysiology of a shiro and thus reveal the mechanism of sporocarp formation. All SNPs, indels, and transposon insertions in the genome were identified, and their chromosomal locations were determined. This information could be used to reveal the genetic diversity of matsutake in nature, conserve its genetic resources, and ensure its production. Furthermore, sequence variant analysis, followed by genome-wide association study, could reveal the genetic mechanisms underlying phenotypic variations in the physiological and metabolomic traits of matsutake. As mentioned above, the matsutake genome assembly constructed in this study could serve as a reference for further genetic studies.

## Data availability

Raw sequence reads were deposited in the Sequence Read Archive (SRA) database of the DNA Data Bank of Japan (DDBJ) under the accession number DRA014434. Assembled sequences are available at DDBJ (accession numbers AP026538 - AP026551) and Plant GARDEN (https://plantgarden.jp).

## Acknowledgments

We thank Prof. S. Kuraku (National Institute of Genetics, Japan) for helpful discussions; T. Kurokochi for providing the matsutake samples; and Y. Kishida, C. Minami, K. Ozawa, H. Tsuruoka, and A. Watanabe (Kazusa DNA Research Institute) for technical assistance. This study was supported in part by JSPS KAKENHI (16K20964, 20H00429, 22H05172, and 22H05181) and the Kazusa DNA Research Institute Foundation.

## Supplementary data

**Supplementary Table S1** Annotation of variants detected among the 14 matsutake lines.

**Supplementary Figure S1** Genomic DNA extracted from dried matsutake sporocarps.

Lanes 1 and 2 indicate the genomic DNA of matsutake samples A and B, respectively. The three molecular weight markers used are as follows: Marker 7 GT (Nippongene, Tokyo, Japan), λ-HindIII digest (Takara Bio, Kusatsu, Japan), and 2.5 kb DNA Ladder (Takara Bio).

**Supplementary Figure S2** Estimation of the genome size of matsutake, based on *k*-mer analysis (*k* = 21) with the given multiplicity values.

**Supplementary Figure S3** Genome coverage of the assemblies generated in previous studies. Blue, green, black, and red lines indicate the genome coverage of Trima3, Tricma30605_assembly01, ASM293902v2, and ASM331463v1, respectively, within a 100 kb window. Gray shadows indicate regions with high GC content.

