## Supplementary Figure for "Telomere-to-telomere genome assembly of matsutake (*Tricholoma matsutake*)"

**Supplementary Table S1** Annotation of variants detected among the 14 matsutake lines.

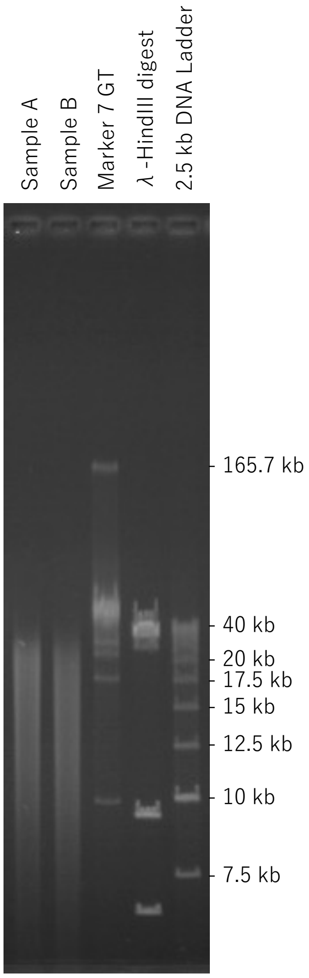


**Supplementary Figure S1** Genomic DNA extracted from dried matsutake sporocarps.
Lanes 1 and 2 indicate the genomic DNA of matsutake samples A and B, respectively. The three molecular weight markers used are as follows: Marker 7 GT (Nippongene, Tokyo, Japan), λ-HindIII digest (Thermo Fisher Scientific, Waltham, MA, USA), and 2.5 kb DNA Ladder (Takara Bio, Kusatsu, Japan).

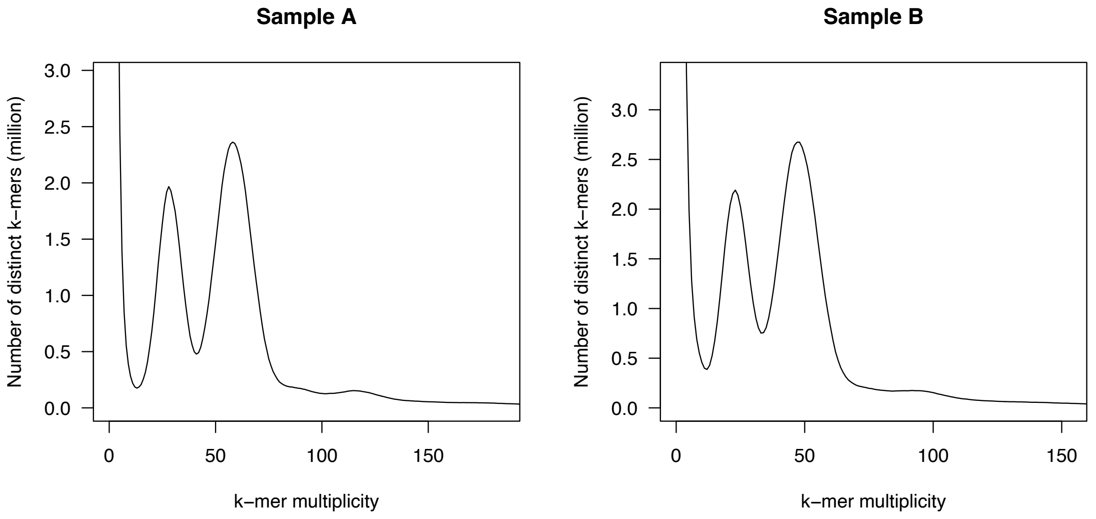


**Supplementary Figure S2.** Estimation of the genome size of matsutake, based on *k*-mer analysis (*k* = 21) with the given multiplicity values.

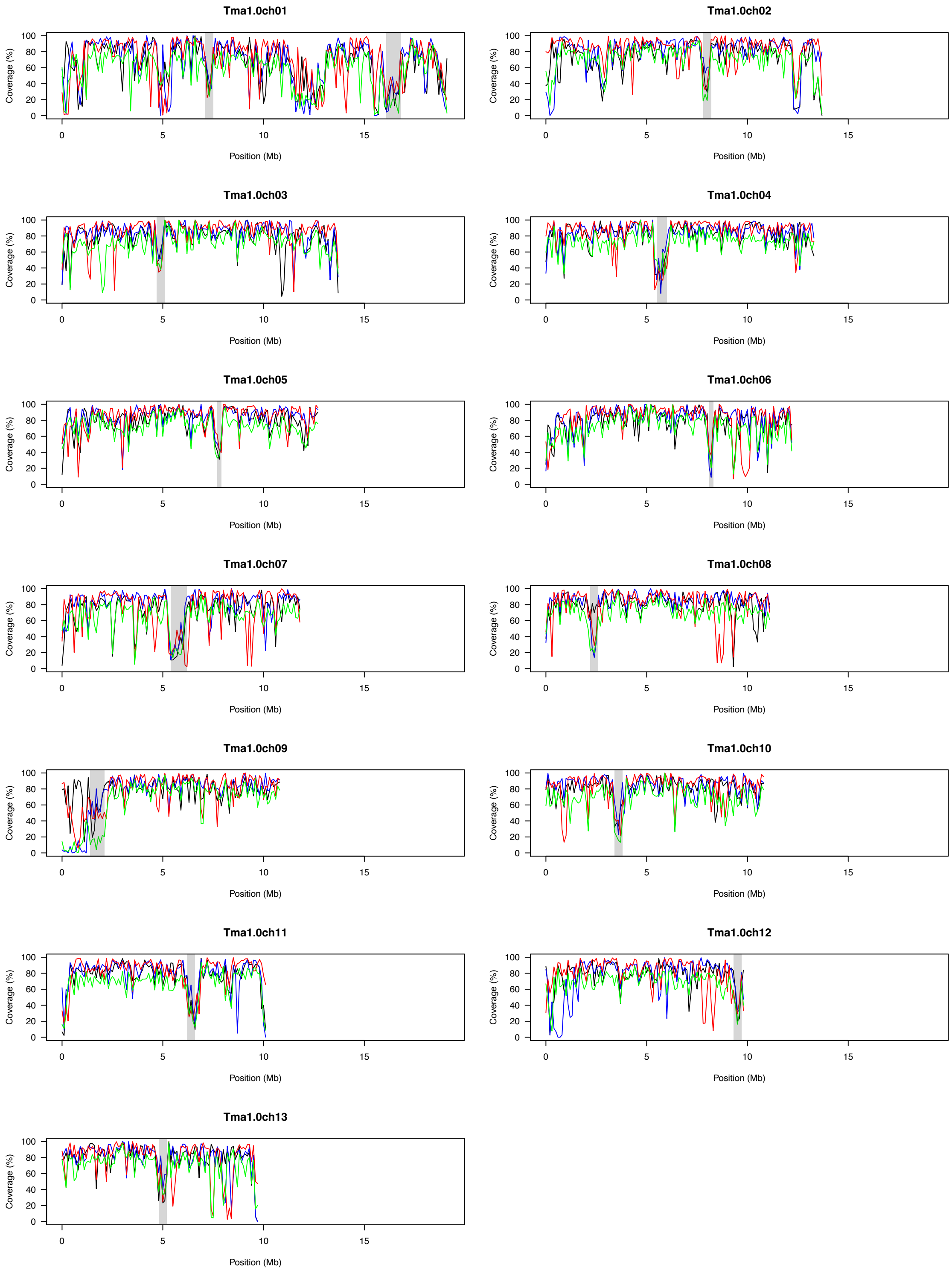


**Supplementary Figure S3** Genome coverage of the assemblies generated in previous studies. Blue, green, black, and red lines indicate the genome coverage of Trima3, Tricma30605_assembly01, ASM293902v2, and ASM331463v1, respectively, within a 100 kb window. Gray shadows indicate regions with high GC content.
